# Specific hippocampal representations are linked to generalized cortical representations in memory

**DOI:** 10.1101/207142

**Authors:** Jai Y. Yu, Daniel F. Liu, Adrianna Loback, Irene Grossrubatscher, Loren M. Frank

## Abstract

Memories link information about specific experiences to more general knowledge that is abstracted from and contextualizes those experiences, but how neuronal activity patterns support this link is not known. Here we show that during memory reactivation in a foraging task with multiple spatial paths, specific hippocampal place representations are concurrently and preferentially reactivated with a subset of prefrontal cortical task representations that generalize across different paths. This link between specific and general representations may serve as a neural substrate for abstraction and task guidance in mammals.

## Main text

How does the brain represent the content of individual experiences with respect to their more general significance? Take, for example, memories of using staircases in your apartment building, home, or workplace. You can most likely recall unique memories of walking up a specific staircase connecting two particular floors. These individual memories do not exist in isolation but are connected to more general knowledge, such as staircases are used for travelling between floors. It is not known how activity patterns in the brain support the link between specific and general features of experience, which is necessary to correctly embed individual memories in broader knowledge structures.

The hippocampus and prefrontal cortex (PFC) are thought to play complementary roles in maintaining these types of knowledge^1–6^. The hippocampus is important for memories of specific experiences^7–9^, while the PFC supports memories that generalize across experiences in the form of “schemas”^10–12^. In the context of rodent spatial exploration, individual experiences involve the activity of hippocampal place cell ensembles that represent specific locations^13–15^ and spatial trajectories^16^. In contrast, PFC firing patterns seen during behavior are highly heterogeneous, and are more often related to different behavioral stages or the structure of an ongoing task^17–29^ than locations in space^18,19^. Hippocampal-cortical representations observed during ongoing experience are transiently reinstated during hippocampal sharp-wave ripple (SWR) memory reactivation^30,31^, which indicates associations observed during ongoing experience are preserved in memory. Given the contrasting activity patterns observed in these regions, it remains unclear how hippocampal-cortical associations reflect the link between representations for specific and general features of experience.

Curiously, only a subset of PFC cells participate in awake SWRs events ^30,32–34^. This suggests that only a select subset of PFC representations remain linked in memory with hippocampal location representations. However, it is not known whether this subset constitutes an unbiased selection from the pool of heterogeneous PFC representations or a selection enriched for certain representations. Determining the properties of these reactivated representations is critical for understanding how hippocampal-cortical activity patterns preserve links between representations for specific and general knowledge.

We therefore examined hippocampal-cortical representations by recording activity simultaneously in the hippocampus and PFC of rats performing a spatial foraging task in which they are required to travel across multiple paths to obtain food reward. The task was designed to examine representations for “specific” and “general” features of experience in a spatial context, where “specific” refers to representations expressed on individual trajectories and “general” refers to representations expressed across multiple trajectories that capture their common features. The task had four potential reward locations (wells) interconnected by paths. At any given time, only two wells were active and could dispense reward. To receive reward the animal needed to find these two wells and visit them in alternation (Fig. S1). By switching the rewarded wells within and/or between sessions or days, we encouraged the animal to travel between wells via different trajectories that can consist of any combination of paths. We defined a trial as the time between consecutive well location visits. To compare hippocampal (CA1) and PFC activity on different trajectories, we normalized CA1 and PFC activity by dividing each trial into 36 time bins (see Methods). We also confirmed the animal displayed similar movement patterns on different trials and trajectories (Fig. S2).

As expected, CA1 cells (n = 234) showed location specific activity (Fig. 1A-C, CA1 cells 1-2). PFC cells (n = 578) showed diverse activity patterns including those that varied across different trajectories (Fig 1A-C, PFC cells 1-2) and those that were active at similar trial phases across different trajectories, consistent with representations of trial structure (Fig 1A-C, PFC cells 3-4). We described the firing pattern of each cell using two parameters: all-trial similarity and maximum within-trajectory similarity. All-trial similarity captures the consistency of firing profiles between all pairs of trials and is thus a measure of generalization across trials. All-trial similarity was quantified as the median of pairwise Pearson’s correlations between firing profiles across all pairs of trials (R_median_, Fig. 1D). Maximum within-trajectory similarity captures the reliability of the cell’s activity in representing aspects of a specific spatial trajectory and was quantified by calculating a similarity score separately for trials on each trajectory and then taking the maximum of those scores across trajectories (R_max_, Fig. 1E).

**Figure 1.**
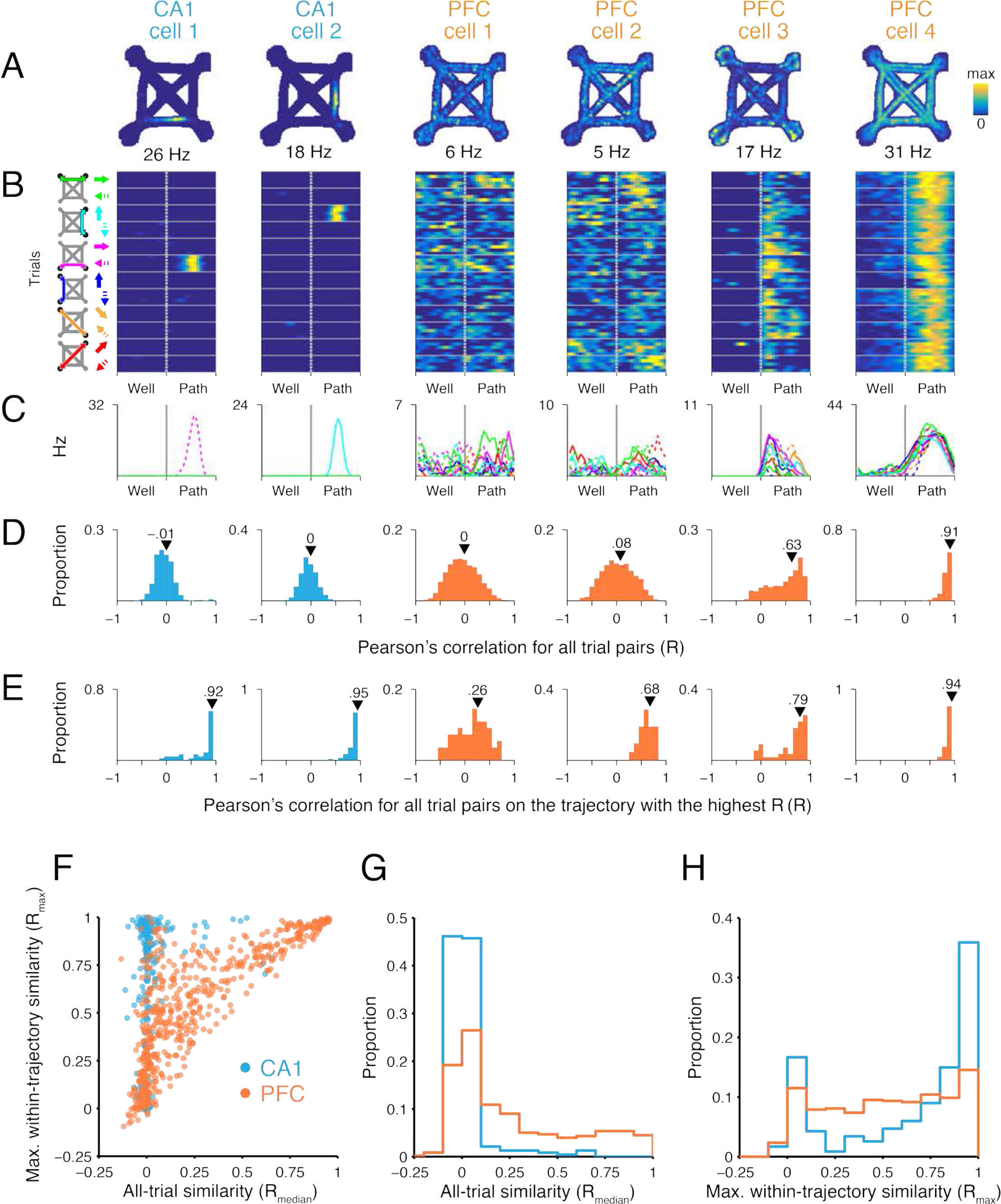
Location specific CA1 and diverse task PFC activity patterns. **A.** Occupancy normalized spatial firing maps for 2 CA1 and 4 PFC cells that were simultaneously recorded. The maximum firing rate for each cell is indicated below each panel. **B.** Time normalized trial firing rate maps for the cells in A. The vertical dotted line separates well and path trial phases. Horizontal solid lines separate trials on each trajectory. 5 example trials are shown for each trajectory. The trajectory for each group of trials is indicated by the schematic on left of column 1. Solid and dotted arrows indicate the two directions of travel on a path. The firing rate color scales are the same as in A. **C.** Median firing rate for each trajectory. Line color corresponds to arrow color scheme in B. **D.** Distribution of pairwise Pearson’s correlation for trial firing profile across all trials. All-trial similarity (R_median_) is the median of the distribution and is indicated by the arrowhead. **E.** Distribution of pairwise Pearson’s correlation for firing profile of trials on the trajectory with the highest median pairwise correlation. Maximum within-trajectory similarity (R_max_) is the median of the distribution and is indicated by the arrowhead. **F**. Scatter of R_max_ and R_median_ for CA1 (cyan, n=234) PFC cells (orange, n=557). **G.** Distribution of R_median_ (from F) for CA1 (cyan, n=234) and PFC cells (orange, n=578). Kolmogorov-Smirnov test: ***p<10^−4^. **H.** Distribution of R_max_ (from F) for CA1 (cyan, n=234) and PFC cells (orange, n=557). Kolmogorov-Smirnov test: ***p<10^−4^.

Across the population, CA1 cells typically had low R_median_ but high R_max_ values (Fig. 1F-H, cyan), reflecting their spatially specific and reliable location representations. In contrast, PFC cells had a wide range of R_median_ and R_max_ values (Fig. 1F-H, orange), reflecting the heterogeneity in trial-related representations (Fig. S3) that is typical for this brain region. Nonetheless, a subset of these PFC cells had similar (high R_median_) and reliable (high R_max_) activity patterns even between trials on trajectories with different lengths (e.g. Fig. 1 PFC cell4 and Fig. S4). The firing properties of these cells are consistent with generalized trial representations irrespective of differences in spatial location and geometry.

We next asked whether a specific subset of these heterogeneous PFC representations are linked with hippocampal spatial representations during awake SWR memory reactivation. Since trajectory representations are frequently reactivated during SWRs^35–37^, our goal was to understand how reactivation of different path-associated representations are coordinated between CA1 and PFC. We therefore selected CA1 and PFC cells that were preferentially active on paths^34^. We then examined SWR events that reactivated these path-active CA1 cells and asked whether during these SWRs, concurrently reactivated PFC cells (i.e. cells with firing rate increases) (n = 35 cells) had task activity patterns that differed from the selected PFC cells that were not reactivated (i.e. cells without firing rate change) (n=161 cells) (Fig. S5).

We found that the population of reactivated PFC cells was substantially enriched for cells with high all-trial similarity (R_median_), which by definition also had high maximum within-trajectory similarity (R_max_)(Fig. 2A-E PFC cells a-b and Fig 2.F-H orange). By contrast, PFC cells that did not participate in SWR reactivations tended to show activity that varied from trial to trial or between trajectories. The activity pattern of this population was mostly dissimilar across trajectories (R_median_ distribution dominated by low values) and varied in within-trajectory similarity (R_max_ values distributed uniformly) (Fig. 2A-E PFC cells c-f and Fig. 2 F-H gray).

**Figure 2.**
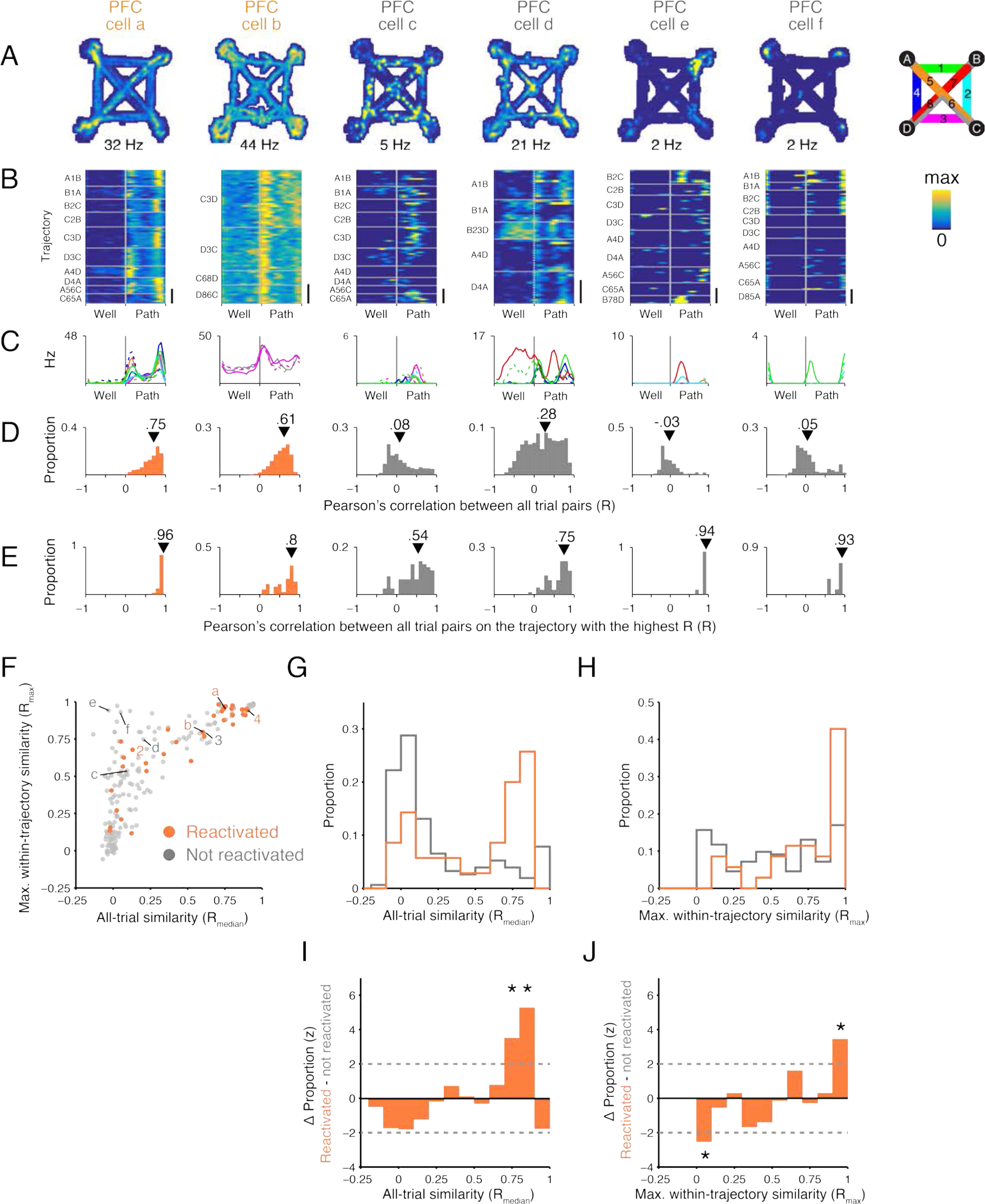
SWR reactivated path-preferring PFC cells show higher activity similarity across different trajectories. **A.** Occupancy normalized spatial firing maps for path-preferring PFC cells. Maximum firing rate of each cell is indicated below each panel. Cells a and b show significant SWR reactivation. Cells c-f are not reactivated. Cells a and c were recorded simultaneously. **B.** Time normalized trial firing rate map for the cells in A. The vertical dotted line separates well and path trial phases. Horizontal solid lines separate trials on each trajectory. Each trajectory is labeled to indicate the start and the end well locations (letters), and paths taken (number) according to the maze schematic in A. Maximum firing rate and color scale are the same as in A. Scale bar: 10 trials. **C.** Median firing rate for each trajectory. Line color key is indicated by maze schematic in A. **D.** Distribution of pairwise Pearson’s correlation for trial firing profile across all trials. All-trial similarity (R_median_) is the median of the distribution and is indicated by the arrowhead. **E.** Distribution of pairwise Pearson’s correlation of trials on the trajectory with the highest median pairwise correlation. Maximum within-trajectory similarity (R_max_) is the median of the distribution and is indicated by the arrowhead. **F**. Scatter of R_max_ and R_median_ for SWR reactivated (orange, n=35) and non-reactivated (n=161, gray) PFC cells. Cells from Fig. 1 (2 and 4) and Fig. 2A (a-f) are labeled. **G.** Distribution of R_median_ (from F). Kolmogorov-Smirnov test: ***p<10^−3^. **H.** Distribution of R_max_ (from F). Kolmogorov-Smirnov test: ***p<10^−3^. **I.** Difference in the R_median_ distributions between path SWR reactivated and not reactivated PFC cells normalized using a permutation test (see Methods). **J.** Difference in the R_max_ distributions between path SWR reactivated and not reactivated PFC cells normalized using a permutation test (see Methods). **I-J.** Dotted lines indicate ±2 S.D․.

The differences between reactivated and non-reactivated PFC cells were clear when we examined the distribution of R_median_ and R_max_ relative to that of the entire path-preferring PFC population. If reactivated PFC cells were an unbiased subsample from the population of path-preferring PFC cells, we would expect to find comparable R_median_ and R_max_ distributions in both the reactivated and non-reactivated populations. Instead, we found a significantly larger fraction of PFC cells with high R_max_ and R_median_ in the SWR reactivated population than expected given the baseline population distribution (Fig. 2 I-J). This difference in SWR engagement could not be explained by differences in peri-SWR firing rate that might have influenced our ability to detect positive SWR modulation in PFC cells (Fig. S6).

Our results demonstrate that SWR reactivation of hippocampal path location representations engages a subset of PFC cells expressing path-related representations that generalize across trajectories (i.e. cells with high all-trial similarity). These findings suggest a many-to-one mapping between hippocampal and PFC representations that is maintained in hippocampal-cortical networks, where many location representations in the hippocampus can be linked to a single generalized representations related to path traversal in the PFC. We tested this prediction by creating groups of SWRs that reactivated different hippocampal location representations, defining each group as the set of SWRs where only one place cell was reactivated. We then compared, across pairs of groups, the firing rate of concurrently reactivated PFC cells, focusing on PFC cells that expressed generalized representations (R_median_>0.5). As predicted, we found SWRs reactivating hippocampal place cells that represented locations on different trajectories could engage the same PFC cell (Fig. 3A).

**Figure 3.**
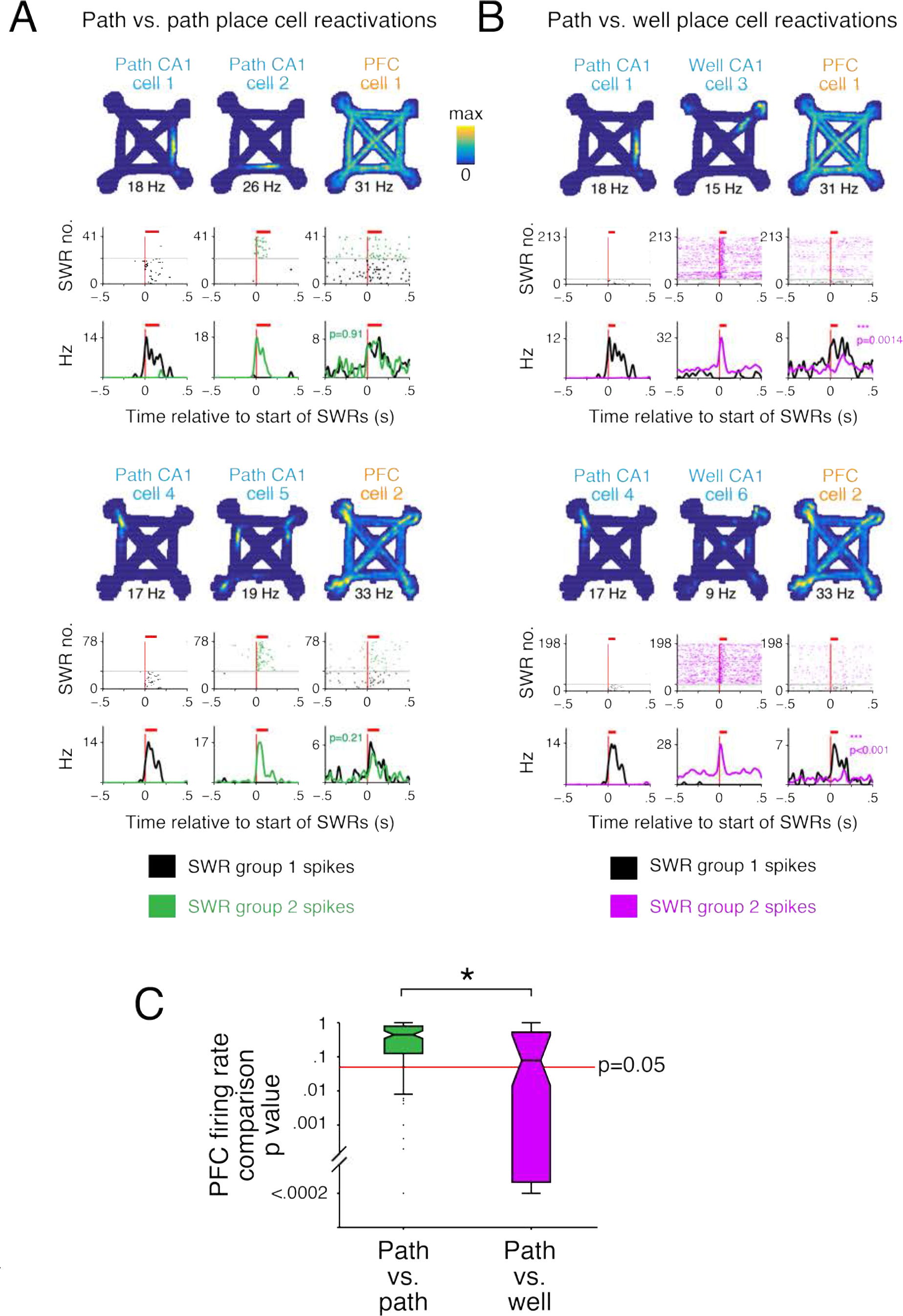
Similar PFC modulation accompanies distinct hippocampal path location reactivations. **A-B.** Spatial firing rate map, SWR aligned spiking raster and firing rate for a pair of CA1 cells and a PFC cell that were recorded simultaneously. Each group of SWRs was defined as events containing only spikes from one of the two CA1 cells (columns 1 and 2). The corresponding spiking and firing rate of the PFC cell during these two groups of SWRs are shown in column 3. The mean duration of SWRs is indicated with a red bar. The difference in PFC firing rates between the two groups of SWRs is quantified using a permutation test. Two such sets are shown (CA1 cells 1-3 and PFC cell 1, and CA1 cells 4-6 and PFC cell 2). In the top set, CA1 cells 1 and 2, and PFC cell 1 corresponds to the CA1 cells and the PFC cell 4 in Fig. 1 respectively. PFC cell 2 in the bottom set corresponds to PFC cell a in Fig. 2. **A.** Comparison of PFC spiking between SWRs containing different path place cell reactivations. **B.** Comparison of PFC spiking between SWRs containing path or well place cell reactivations. **C.** Boxplot of permutation test p values for comparisons of PFC firing rates during SWRs between groups of SWRs. PFC cells with similar firing patterns across trajectories (R_median_>0.5) were included. PFC activity during SWRs reactivating different CA1 path location representations was similar (green, higher p values, n=127 SWR group pairs, examples in A) compared with activity during SWRs reactivating CA1 path or well location representations was dissimilar (magenta, lower p values, n=87 pairs, examples in B). Wilcoxon rank-sum test: ***p<10^−3^.

Importantly, this many-to-one mapping reflected task relevant associations, and was not a result of non-specific activation of PFC cells across all SWRs. Here we took advantage of our recent demonstration that the reactivation of hippocampal representations for locations associated with movement (i.e. paths) is largely distinct from those for locations of immobility (i.e. wells)^34^. Based on that observation, we reasoned that generalized PFC path representations should be reactivated together with hippocampal path location representations but not with hippocampal well location representations.

Consistent with this prediction, we found PFC activity during SWRs was often different between SWRs defined by the reactivation of either a path or a well location active CA1 cell (Fig. 3B). Comparisons of PFC firing rates during these groups of SWRs showed more significant differences than for comparisons between groups of path versus path place cell SWRs (see methods) (Fig. 3C). This indicates that PFC reactivation patterns distinguished between path versus well location reactivations, whereas different hippocampal path location reactivations were often associated with similar PFC reactivation patterns. These results confirm that the many-to-one mapping between hippocampal and cortical activity patterns reflected task information and is consistent with our previous finding of coordinated hippocampal-cortical reactivation during awake SWRs^30,34^.

Our results showed that during SWR memory reactivation, generalized path representations in PFC are preferentially and coherently linked with specific location representations in CA1. We suggest the enrichment of these links reflects the network’s ability to associate representations for frequently repeating features of ongoing experience (i.e. the repeating structure of each trial) with features specific to an experience (i.e. traversal of one particular trajectory). Enriching general-to-specific links in the network could provide a basis for creating abstractions based on similar experiences as an animal learns^4–6^, which requires the function of hippocampal-cortical networks^1–4,7,8,10,38–46^. These abstractions could facilitate the creation of “schemas” or generalizable knowledge of a task^47,48,49^ that are crucial for guiding future decisions^11,12^.

## Author contributions

J.Y.Y and L.M.F. designed the experiments and wrote the manuscript. J.Y.Y. analyzed the data and performed experiments for the foraging task with assistance from I.G., D.L. and A.L.

## Acknowledgments

We thank B. Mensch, G. Rothschild, Mari Sosa, E. Anderson, D. Kastner, E. Phillips, C. Geaghan-Breiner, A. Gillespie, H. Joo and A. Kiseleva for comments on the manuscript. This work was supported by a Jane Coffin Childs Memorial Fund for Biomedical Research postdoctoral fellowship (J.Y.Y.), the Howard Hughes Medical Institute, NIH R01MH105174, NIH R01MH097084 and University of California Office of the President Lab Fees Award #LF-12-237680 (L.M.F.).

## Data availability

The data that supports the findings of this study is available in the Collaborative Research in Computational Neuroscience data repository (http://dx.doi.org/10.6080/K0NK3BZJ).

## Methods

### Animal and behavior

All experiments were conducted in accordance with University of California San Francisco Institutional Animal Care and Use Committee and US National Institutes of Health guidelines. We trained 6 rats initially to traverse a 1m long linear track for reward (evaporated milk plus 5% sucrose, Carnation). We then introduced the animals to the foraging task ~21 days after surgery. In the task, only two of the four possible reward well locations were chosen to deliver reward. The rat had to find those two well locations and visit them in alternation to receive reward. We changed the rewarded well locations within or between sessions, or between days. These changes in spatial reward contingencies were not explicitly signaled and the rat needed to find the new rewarded well locations by trial and error. Data from 11, 9, 10, 12, 13 and 4 days were analyzed for each of the 6 animals respectively. Each day consisted of between 2-3 task sessions of 15-45 minutes each. The task sessions were interleaved with rest sessions of 20-60 minutes in a sleep box located away from the track. We used a custom built automated system for reward delivery, which was triggered by an infrared beam break at the well location. Reward was delivered immediately (2 animals) or after a 1s delay (4 animals) after the beam break. A syringe pump (NE-500 OEM, New Era Pump Systems Inc.) delivered the reward (100-300μ1 at 20ml/min).

### Implant

Custom designed and 3D printed (PolyJetHD Blue, Stratasys Ltd.) recording drives housed a maximum of 28 individually movable tetrodes. Tetrodes (Ni-Cr, California Fine Wire Company) were gold plated to reach an impedance of 250kOhm at 1kHz.

Implanted recording drives targeted both dorsal CA1 (7 tetrodes) and dorsal PFC (14-21 tetrodes, housed in one cannula angled at 20 degrees toward the midline). CA1 AP: -3.8mm and ML: 2.2mm. PFC (anterior cingulate cortex and dorsal prelimbic cortex): AP: +2.2mm, ML +1.5mm and DV between 1.88mm to 2.72mm depending on the AP and ML coordinates of each tetrode.

Tetrodes were adjusted every 2 days post-surgery to reach the target DV coordinate (PFC) or guided by LFP and spiking patterns (CA1). After the start of data acquisition, tetrodes were adjusted at the end of the day in small increments (typically ~30μm) to improve cell isolation.

### Histology

At the end of the experiment, we marked the location of recording sites by passing current through each tetrode (30μA, 3s) to create electrolytic lesions. After 12-24 hours, we perfused the animals with paraformaldehyde (4% in PBS), fixed (24 hours at room temperature) and cryoprotected the brain (30% sucrose in PBS at 4°C). We identified the sites of electrolytic lesions with Cresyl Violet stained coronal sections (50μm).

### Recording

The NSpike data acquisition system (LMF and J. MacArthur, Harvard Instrumentation Design Laboratory) was used for data collection. Experiments were conducted in dim lighting. To track the animal’s position, an infrared LED array was mounted on the headstage amplifier and video was recorded at 30Hz. We recorded LFP from each tetrode (0.5-400Hz sampled at 1.5kHz). We recoded spiking activity from each tetrode channel (600-6000Hz or 300-6000Hz sampled at 30kHz). For hippocampal tetrodes, the reference for LFP and spike detection was a tetrode located in corpus callosum. For PFC, the reference was a tetrode located locally but did not detect spikes.

### Data analysis

We identified putative neurons by manual clustering of spiking data from channels of each tetrode based on peak amplitude, spike width and wave-form principal components (MatClust, M.P.K.). Only stable and well-isolated cells were used for further analysis.

The animal’s position was determined as the centroid of the front and back diodes from the LED array using a semiautomated analysis of the video.

### Cell selection

CA1: We recorded from 391 CA1 neurons from which we excluded from our analysis putative fast spiking interneurons (spike peak to trough width <0.4ms and mean firing rate >10Hz, n=22) and cells with <200 spikes across all sessions on a given day (n=135).

PFC: We recorded from 844 PFC from which we excluded from our analysis putative fast spiking interneurons (spike peak to trough width <0.3ms and mean firing rate >7Hz, n=42) and cells that had <200 spikes across all sessions of a day (n=224). We defined path active PFC cells as those with peak firing during the path phases of the trial.

### SWR detection

The raw CA1 LFP was referenced to an electrode in corpus callosum and then filtered (150-250Hz) to isolate the SWR band. The SWR envelope was then obtained using the Hilbert transform and convolved with a Gaussian kernel (σ=4ms). A consensus SWR envelope was calculated by taking the median of the envelopes across all available tetrodes. Only days where at least three tetrodes were in or near the CA1 cell layer were used for the analysis.

To avoid the problem of arbitrary thresholds and differences in noise distributions across days, we developed a SWR identification approach that defines a threshold based on the distribution of the consensus envelope power for a given day^34^. Only SWR events occurring at speed <4cm/s were included in our analyses.

### Occupancy normalized firing maps

The environment was first divided into 2cm square bins. The occupancy-normalized rate was calculated by dividing the number of spikes by the occupancy of the animal per bin and smoothing with a 2-dimensional symmetric Gaussian kernel (σ=2cm and 12cm spatial extent).

### Trial normalization

Each trial was defined as the time between entry to a well location and entry to the next well location. For each trial, the time the rat spent at the well location was divided into 18 equally spaced time bins. The same was done to time when the rat was travelling between well locations. Thus each trial comprises of 36 bins (Fig. 1B and Fig. 2B). Firing rate for each bin was calculated by dividing the number of spikes that occurred during each time bin by the duration of each time bin. The firing rate was smoothed using a Gaussian kernel (σ=1 trial bin). Time and spiking during SWRs and within +/− 50ms were excluded and does not contribute towards the calculation of firing properties (e.g. R_median_ and R_max_). The speed for each bin is the mean of speed of time points in each bin. The speed profile for each trial was smoothed using a Gaussian kernel (σ=1 trial bin).

### Trial firing and speed profile similarity

All-trial firing similarity for each cell is the median of Pearson’s correlation of firing profile between all trials (R_median_, Fig. 1D). The same procedure was used to calculate trial speed profile similarity. To calculate maximum within-trajectory similarity, we first computed the median of pairwise Pearson’s correlation between pairs of trials on a trajectory and repeated this for all trajectories. We then selected the maximum out of these values (R_max_, Fig. 1E).

We generated a control for the effect of trial phase related firing on the similarity measure by circularly permuting the firing profile for each trial, which preserves the relationship between consecutive task phase bins for each trial but disrupts consistency in task phase relationship across trials (Fig. S3).

To ensure the animal’s movement pattern was stereotyped across trials, both during times when the rat was at the reward location and on the path, we only included rewarded trials where the animal’s probability of alternation was >75, which was estimated using a state-space model^50^. In addition, to ensure the reliability of the firing pattern of a cell on each trajectory was adequately sampled, only trajectories on which the animal made 5 or more traversals were included in the analyses. All-trial similarity and maximum within-trajectory reliability were only reported for cells with spiking during trials that fulfilled these selection criteria.

### PFC SWR modulation index

The significance of modulation was calculated as describe previously^30,34,51^. We first generated a perievent time histogram (PETH) for all events aligned to the start of SWRs for the observed data. We then generated a control dataset by circularly permuting the spike times for each SWR event, such that all spikes around one SWR event were circularly shifted by the same amount but this amount varied between SWR events. From this control dataset we then generated a PETH. This was repeated 1000 times. Next we calculated the squared deviation of the observed PETH from the mean of the 1000 control PETHs for the average duration of SWRs for the given type of SWR. We then compared the squared deviation of each of the 1000 control PETHs to the mean of all 1000 control PETHs. The significance value was the fraction of 1000 control PETH deviations that are larger than the observed PETH deviation. We defined SWR reactivated path active PFC cells as those with a significant excitation during SWRs containing CA1 path place cells.

### Permutation test for distribution comparisons

To normalize the observed differences in the R_median_ (Fig. 2I) and R_max_ (Fig. 2J) distributions between SWR reactivated and non-reactivated PFC populations, we used a permutation test to generate expected distributions of differences. First, we permuted the identities of SWR reactivated and non-reactivated PFC cells and generated their corresponding probability density functions (PDFs). We then calculated the difference between the PDFs of the two groups of the permuted dataset. We calculated the mean and standard deviation for each bin of the PDF using 10000 permuted datasets with which we used to convert the observed data into z-scores.

### Resample method for matching peri-SWR firing rates

We controlled for the possibility that differences in peri-SWR firing rate could influence our ability to detect SWR modulation and give rise to the observed differences in all-trial similarity (R_median_) between reactivated versus non-reactivated groups. This was achieved by resampling SWR reactivated and non-reactivated PFC cells to match their peri-SWR firing rates (Wilcoxon rank-sum test p values >0.05) (Fig. S6A). The peri-SWR firing rate is the mean firing rate in a 1 second window centered on the start of SWRs. For each resampled dataset, we then obtained the p value of the Wilcoxon rank-sum test for the corresponding all-trial similarity comparison (Fig. S6B). This was repeated 1000 times. The proportion of the 1000 resamples with R_median_ p<0.05 was tested against the expected proportion of resamples with p<0.05 (5%) using a Binomial test (Fig. S6 C-D).

### PFC modulation between different SWR groups

We compared the mean firing rate of two groups of unique SWRs, each containing a different hippocampal place cell. The place cells were active on paths (peak rate >3Hz) or active at a reward well location^34^ (mean rate>3Hz). We extended this to all available pairs of place cells. The mean rate of a PFC cell during each group of SWRs was calculated by dividing the number of spikes observed in each SWR by the mean duration of all SWRs in both groups of a pairwise comparison. A permutation test (5000 permutations) was used to determine if the mean rate of the PFC cell is significantly different between the two groups of SWRs.

---

**Figure S1.**
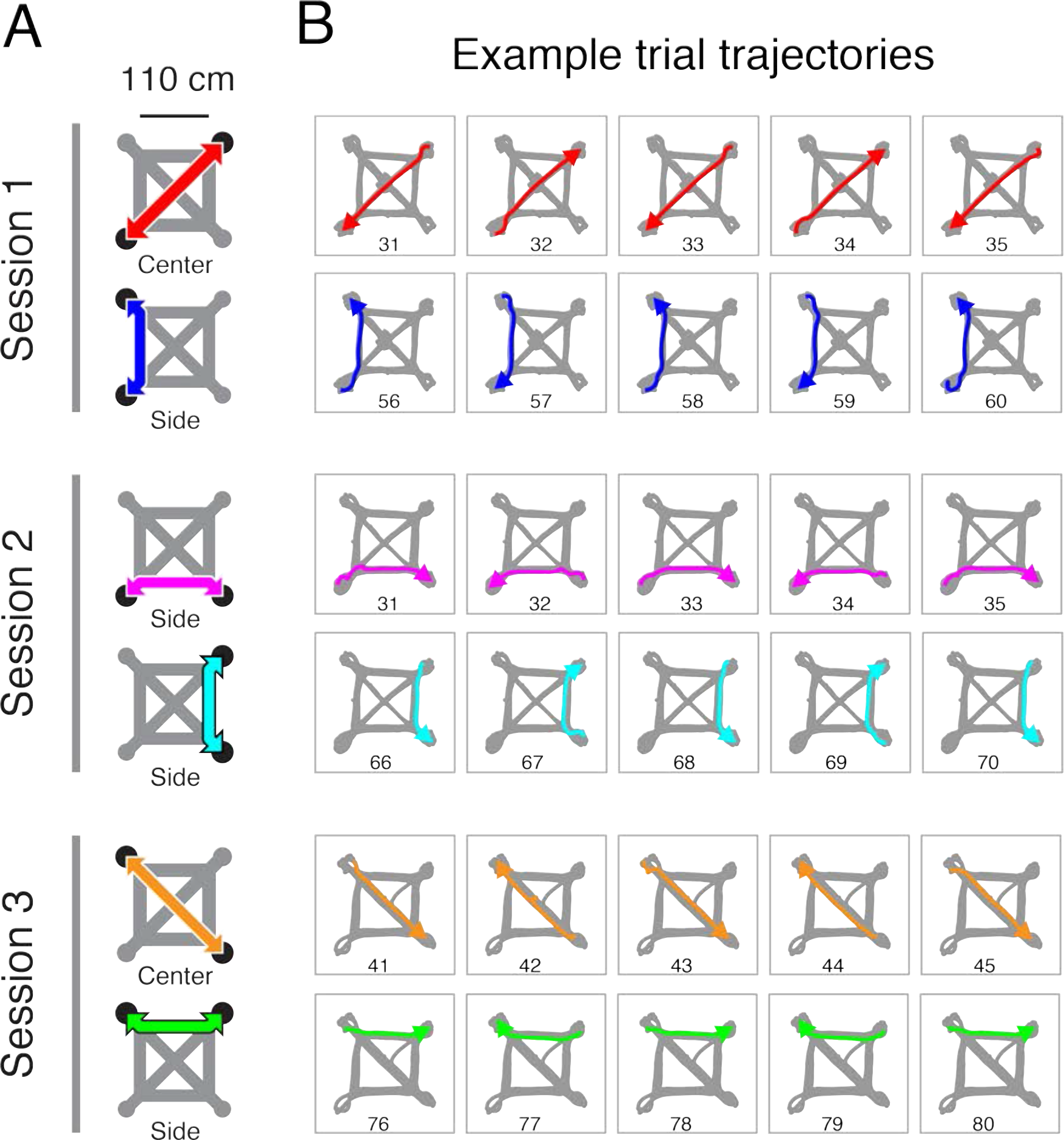
Task structure and example trajectories. **A.** Arrangement of rewarded well locations (black circles) and the most direct interconnecting path (colored arrows) in the maze (gray). The type of path is indicated below each schematic. “Center” refers to paths that cross the center of the maze. “Side” refers to paths on the sides of the maze. The reward well locations changed within and between sessions. **B.** The rat’s trajectory on 5 consecutive trials for each reward well location contingency. The destination well is indicated by an arrow. The trial number relative to the start of each session is indicated.

**Figure S2.**
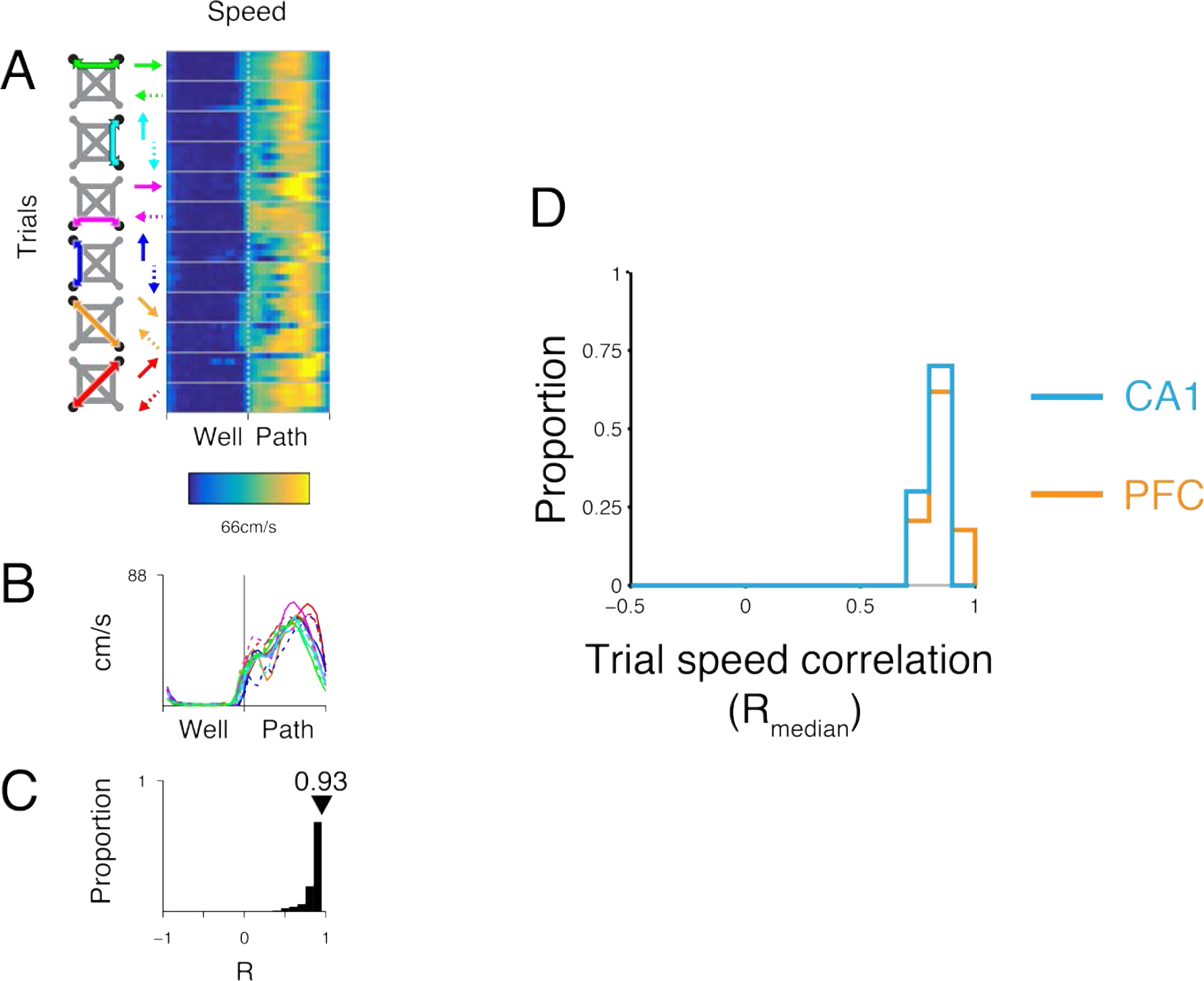
Speed profile similarity. **A.** Trial speed profile for trials shown in Fig. 1. The trajectory of each trial is indicated by the schematic on left. Solid and dotted arrows indicate opposing directions of travel on a path. **B.** Median speed for each trajectory. Line color key correspond to arrow color scheme in A. **C.** Distribution of Pearson’s R for pairwise trial speed profile correlations. The median of the distribution (R_median_) is indicated in red. **D.** Trial speed profile R_median_ for trials on days used for calculating CA1 (cyan, n=10) and PFC (orange, n=35) firing similarity (Fig. 1E). Kolmogorov-Smirnov test: not significant.

**Figure S3.**
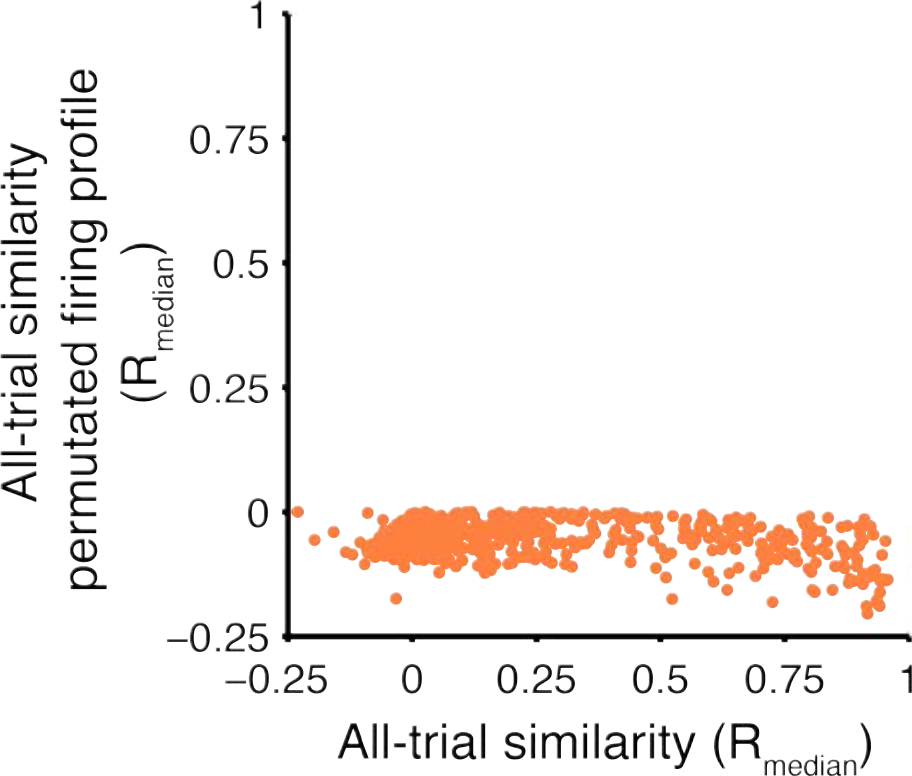
R_median_ is dependent on trial phase. Scatter of R_median_ against mean of R_median_ values for circularly permuted trial firing profiles (1000 permutations for each cell).

**Figure S4.**
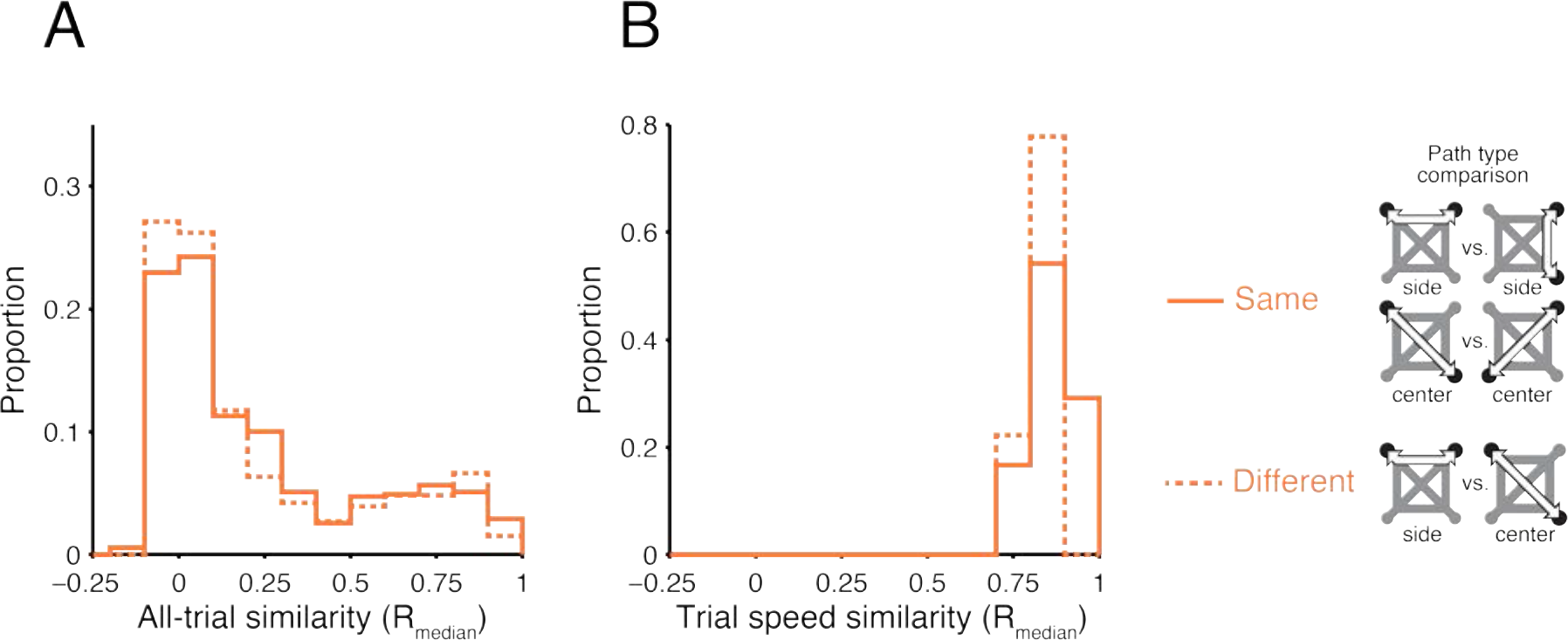
PFC firing is similar within and across trajectories with different lengths. **A.** R_median_ distribution for different path types for PFC units (same n=549 and different n=332). Kolmogorov-Smirnov and Wilcoxon rank-sum tests: not significant. **B.** R_median_ distribution of trial speed similarity for days used to calculate firing similarity in A (same n=24 and different n=9). Kolmogorov-Smirnov and Wilcoxon rank-sum tests: not significant. “Same” indicates comparisons between side with side or center with center trials. “Different” indicates comparisons between side and center trials.

**Figure S5.**
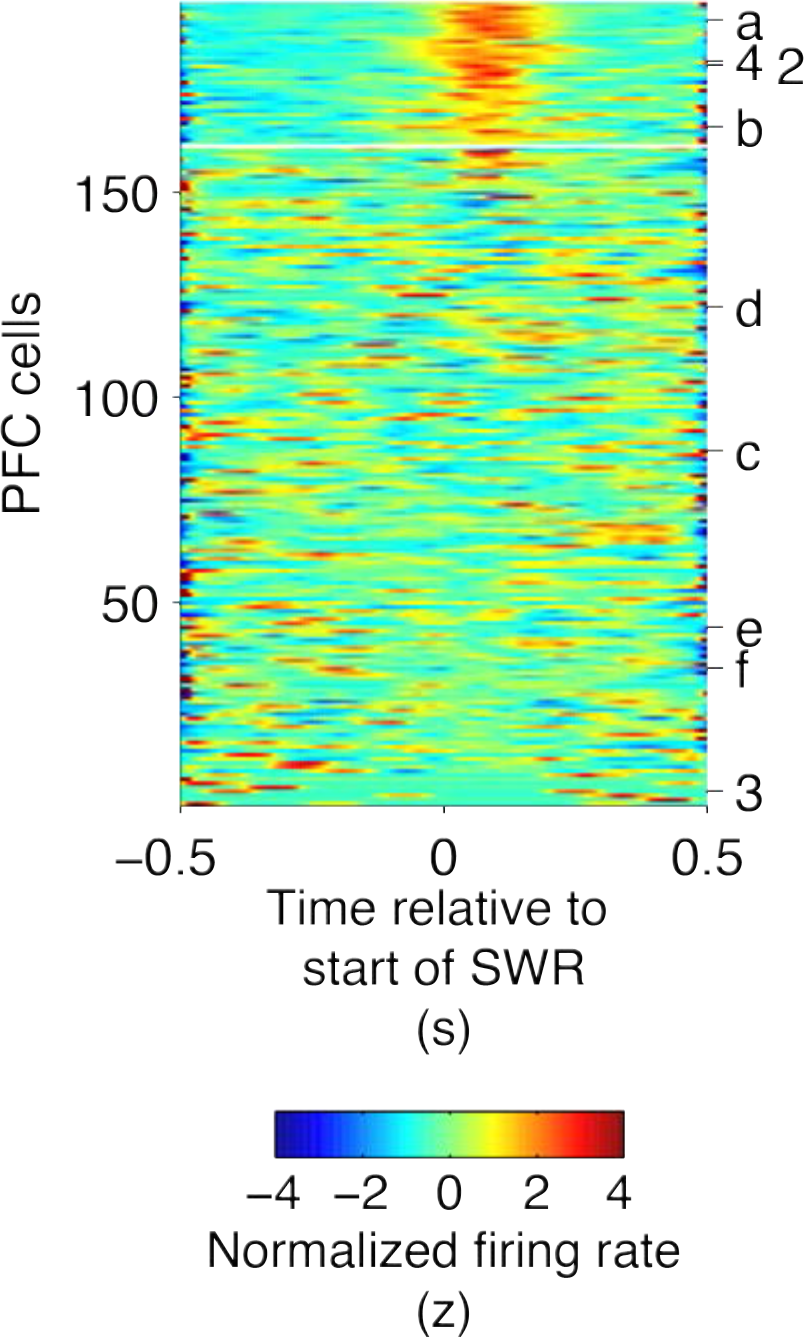
SWR aligned firing of PFC path-preferring cells. SWR aligned normalized firing rate for path-preferring PFC cells. Rows sorted by SWR modulation significance. Cells above the white line show significantly excitation (p<0.05) during path location SWRs. Example PFC cells from Fig. 1 2-4 and Fig. 2 a-f are indicated.

**Figure S6.**
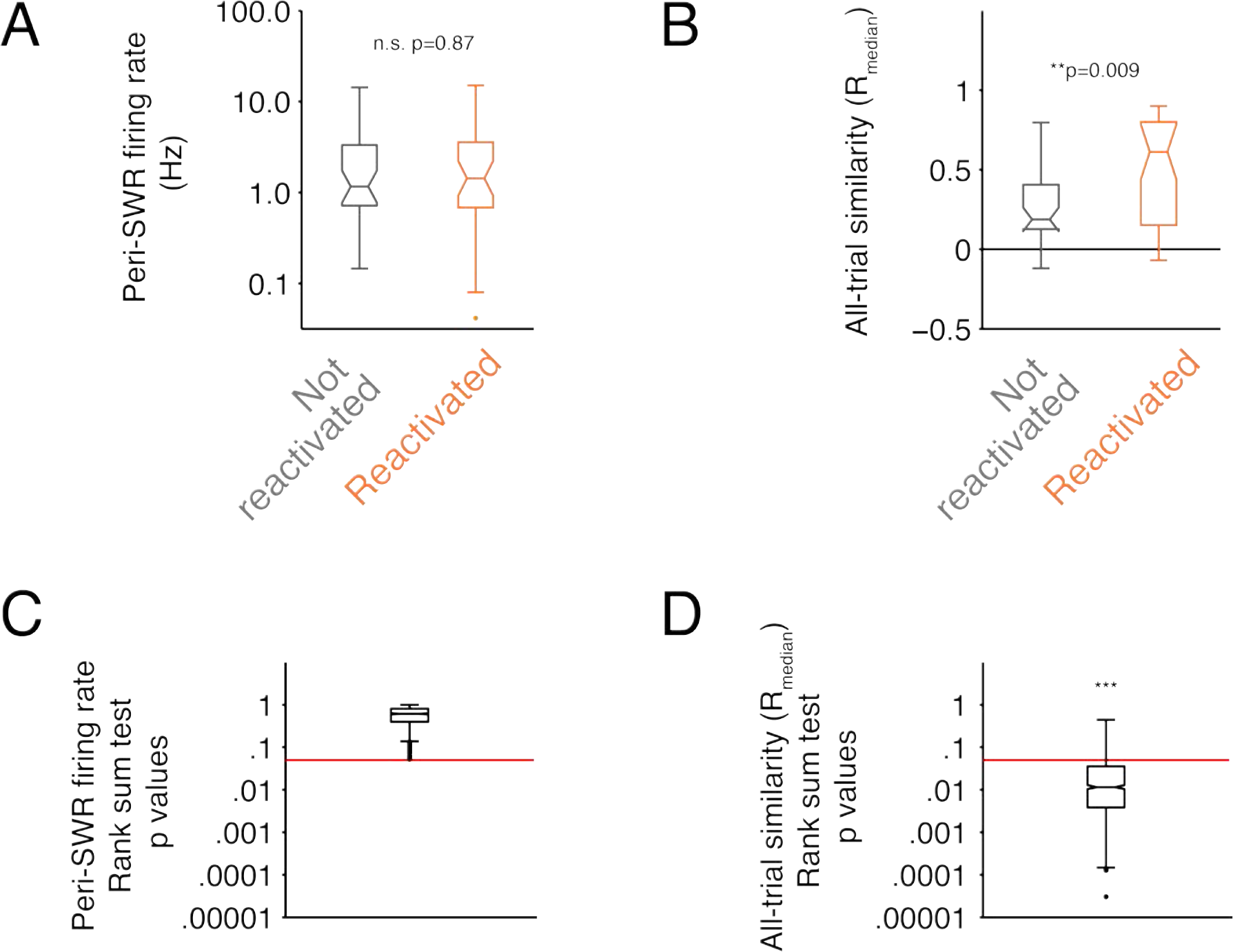
Higher all-trial similarity of SWR reactivated PFC cells compared with non-reactivated cells remains after matching peri-SWR firing rates. **A.** Boxplot of peri-SWR firing rates from an example resampled dataset where PFC cells from the non-reactivated group (gray) were selected to match the peri-SWR firing rates and the number of cells (n=35) from the reactivated group (orange). **B.** Boxplot of corresponding all-trial firing similarity (R_median_) for resampled data in A. SWR reactivated cells (orange) show higher all-trial similarity compared with non-reactivated cells (gray) after matching peri-SWR firing rate. **C.** Boxplot of Wilcoxon rank sum test p values for 1000 peri-SWR firing rate matched resamples. Red line indicates p=0.05. **D.** Boxplot of Wilcoxon rank-sum test p values for differences in all-trial similarity for the corresponding 1000 resamples in B. Binomial test for observed vs. expected proportion of resampled datasets with a significant difference (p<0.05, 5%) is ***p<10^−4^. Red line indicates p=0.05.

